# Role of the amino acid mutations in the HA gene of H9N2 avian influenza virus under selective pressure in escape vaccine antibodies

**DOI:** 10.1101/2021.09.07.459361

**Authors:** Rui Zhu, Shunshun Xu, Wangyangji Sun, Quan Li, Huoying Shi, Shifeng Wang, Xiufan Liu

## Abstract

It has been well-documented that some amino acid mutations in hemagglutinin (HA) of H9N2 avian influenza virus (H9N2 virus) alter the viral antigenicity, but little is reported about the role of antibody escape mutations in escape vaccine antibodies. In this study, we found that the evolution of F/98 strain in chicken embryos or chickens resulted in significant differences in immune escape, and identify the contribution of HA mutations to the antigenic variation and immune escape of H9N2 virus. Among amino acid mutations in the HA of the antigen variant viruses occurring in embryonated chicken eggs and/or chickens with or without the selection pressure of vaccine antibodies, the mutations, S145N, Q164L, A168T, A198V, M224K and Q234L, affect the antigen drift of H9N2 virus. Specially, the A198V mutation, located at the receptor-binding site on the head domain of HA, significantly contributed the antigenic variation of H9N2 virus. The mutation A198V or Q234L significantly improved the receptor binding activity, while S145N mutation decreased the receptor binding activity. Single S145N mutation could promote viral escape from polyclonal antibodies (pAbs) by preventing Ab binding physically, and single A198V mutation could promote viral escape from pAbs by enhancing the receptor binding activity. Additionally, either the mutation S145N or A198V did interfere with the immunogenicity of the inactivated vaccine, resulting in reduction of the protective efficiency of H9N2 inactivated vaccine, which contributed escape from the antibody-based immunity. Our findings provided an important reference for the accurate evaluation of the role of the amino acids mutation in HA affecting the antigenicity of H9N2 virus on immune escape, and delivered a new perspective for monitoring the adaptive evolution of H9N2 virus.

**Importance:** In this study, the role of the HA mutations of H9N2 virus occurring with and without antibody selective pressure on escaping from the antibody-based immune response in host was analyzed. The results demonstrated that (i) the HA mutations S145N, Q164L, A168T, A198V, M224K, and Q234L occurring in the process of the adaptive evolution of H9N2 virus in embryonated chicken eggs and/or chickens could affect the antigenic variation of H9N2 virus. Among these mutations, the HA mutation A198V had the most significant effect on the antigenic variation; (ii) S145N mutation promoted viral escape from pAbs by preventing Abs binding physically; (iii) A198V mutation did promote viral escape from pAbs by enhancing the receptor binding activity; (iv) neither the HA mutation S145N or A198V interfered with the immunogenicity of the inactivated vaccine, resulting in reduction of the protective efficiency of H9N2 inactivated vaccine.

## Introduction

H9N2 avian influenza virus (AIV) spread rapidly and infected more than 90% of chicken flocks since its breakout in Hebei province, China in 1998. It became one of the most important epidemics in poultry industry in China (1). Since then, vaccination strategy of inactivated vaccine for control of H9N2 avian influenza had been extensively executed, and worked well for a long period (2). However, H9N2 virus is undergoing adaptive evolution under the vaccine immune pressure. As a major antigen and receptor binding protein of H9N2 virus, the haemagglutinin (HA) from the circulating field strains were clustered into three lineages before 2007, A/Chicken/Beijing/1/94-like (BJ/94-like), A/Quail/Hong Kong/G1/97-like (G1-like), and A/Duck/Hong Kong/Y439/97-like (Y439/97-like) (3). In 2013, G57 strains were emerged as a predominant genotype of H9N2 virus. A new genotype G118 was discovered in 2015 (4). With the evolution of H9N2 virus, the specific antibodies induced by inactivated vaccine could not effectively block the attachment of HA of the circulating virus to the target cells (5). This resulted in the decrease in the protection efficacy of the existing vaccines and isolation of breakthrough H9N2 viruses isolated in vaccinated chicken flocks with high antibody titer (6). Therefore, it is important to monitor antigenic mutation of the HA of H9N2 virus.

Currently, over 30 antigenic sites of H9N2 virus have been reported (7-13). In these studies, most of mutations on the antigenic sites of HA were gained by monoclonal antibody (mAb) precisely, which promoted virus to escape from mAb neutralization by changing virus-Ab binding. Naturally, the occurrence of antigenic drift of HA is determined by several factors, such as environment, genetic background, and immune status. The immune selective pressure and natural selection could also drive HA mutation on antigenic sites to escape neutralizing antibody by adding N-linked glycosylation (NLG) for shielding the antigenic sites (14, 15), changing virus-antibody binding property (16), or altering receptor-binding specificity (17-20).

We previously reported that the H9N2 vaccine representative strain A/Chicken/Shanghai/F/1998 (F/98, H9N2), which belonged to BJ/94-like lineage, occurred antigenic variation continually when passaged in specific pathogen-free (SPF) chicken embryos or SPF chickens with or without homologous vaccine antibodies (21, 22). In order to identify the contribution of HA mutations to the antigenic variation and immune escape of H9N2 virus, recombinant viruses containing single HA mutation or multiple HA mutations which might affect the antigenic variants of F/98 strains in F/98 backbone were generated to define the role of HA mutations of F/98 strain passaged under the selection pressure with or without homologous vaccine antibodies in immune escape. We found that the evolution of F/98 strain in chicken embryos or chickens resulted in significant differences in immune escape. The results showed that the HA mutations under the selection pressure with or without vaccine antibodies, including S145N, Q164L, A168T, M224K and Q234L, had some effect on the antigenic drift of H9N2 virus, and the HA mutation A198V significantly affected the antigenic variation. Although the virus possessing the HA mutation S145N or A198V escaped from the pAbs-neutralization reaction, the molecular mechanism of antibody neutralization was different between the mutations S145N and A198V, S145N mutation promoted viral escape from pAbs by preventing Abs binding physically, whereas A198V mutation by enhancing the receptor binding activity. Additionally, each of the mutations S145N and A198V interfered with the immunogenicity of the inactivated vaccine, resulting in reduction of the protective efficiency of H9N2 inactivated vaccine, which contributed escape from the antibody-based immunity.

## Results

### The adaptive evolution of H9N2 virus in SPF embryonated chicken eggs or SPF chickens drove different immune escape

We previously reported that antigenic drift of the passaged virus occurred when the F/98 strain passaged continuously in the 47^th^ generation under selective pressure from vaccine antibodies and in the 52^nd^ generation without selective pressure from vaccine antibodies in SPF embryonated chicken eggs (21). The second-generation quasispecies of F/98 strain under selection pressure from vaccine antibodies had undergone 100% antigenic variation in SPF chickens, while after passaging to the fifth generation without selection pressure from vaccine antibodies, only 30-40% of the quasispecies displayed antigen drift (22). The antigenic variant evF47, the 47^th^ generation under vaccine antibodies in SPF chicken embryonated eggs, has mutations K131R, S145N, G181E and A198V in HA. The antigenic variant enF52, the 52^nd^ SPF embryonated chicken eggs SPF embryonated chicken eggs generation without vaccine antibodies in SPF embryonated chicken eggs, also has same mutations K131R, S145N, G181E and A198V in HA. The antigenic variant cvF20, the 20^th^ generation with vaccine antibodies in SPF chicken, has mutations K131R, A198V and Q234L in HA. The antigenic variant cnF20, the 20^th^ generation without vaccine antibodies in SPF chicken, has mutations A168T, A198V and M224K in HA. The above 4 strains were the first antigenic variants, whose genome were still stable after 3 generations of embryos blind passage, and used to determine the role of HA mutations in escaping from antibody-based immune responses with different selection pressures or different models with F/98 strain as a control. SPF chickens immunized by oil-emulsion of inactive whole virus vaccine of F/98 strain were challenged with each of antigenic variants. The virus shedding was detected in tracheal swabs from chickens to analyze the differences in immune escape. The results showed that 100% of SPF chickens (6/6) immunized with F/98 vaccine shed viruses at 3 days after challenge with the virus evF47 or enF52; 66.7% of chickens (4/6) shed virus cvF20; 16.7% of chickens (1/6) shed virus cnF20 (Figure 1). These results revealed that the adaptive evolution of H9N2 virus between embryonated chicken eggs and chickens drove different immune escape, and indicated that the contribution of HA mutations from the above antigenic variants to antigenic drift and immune escape might also be different.

**Figure 1.**
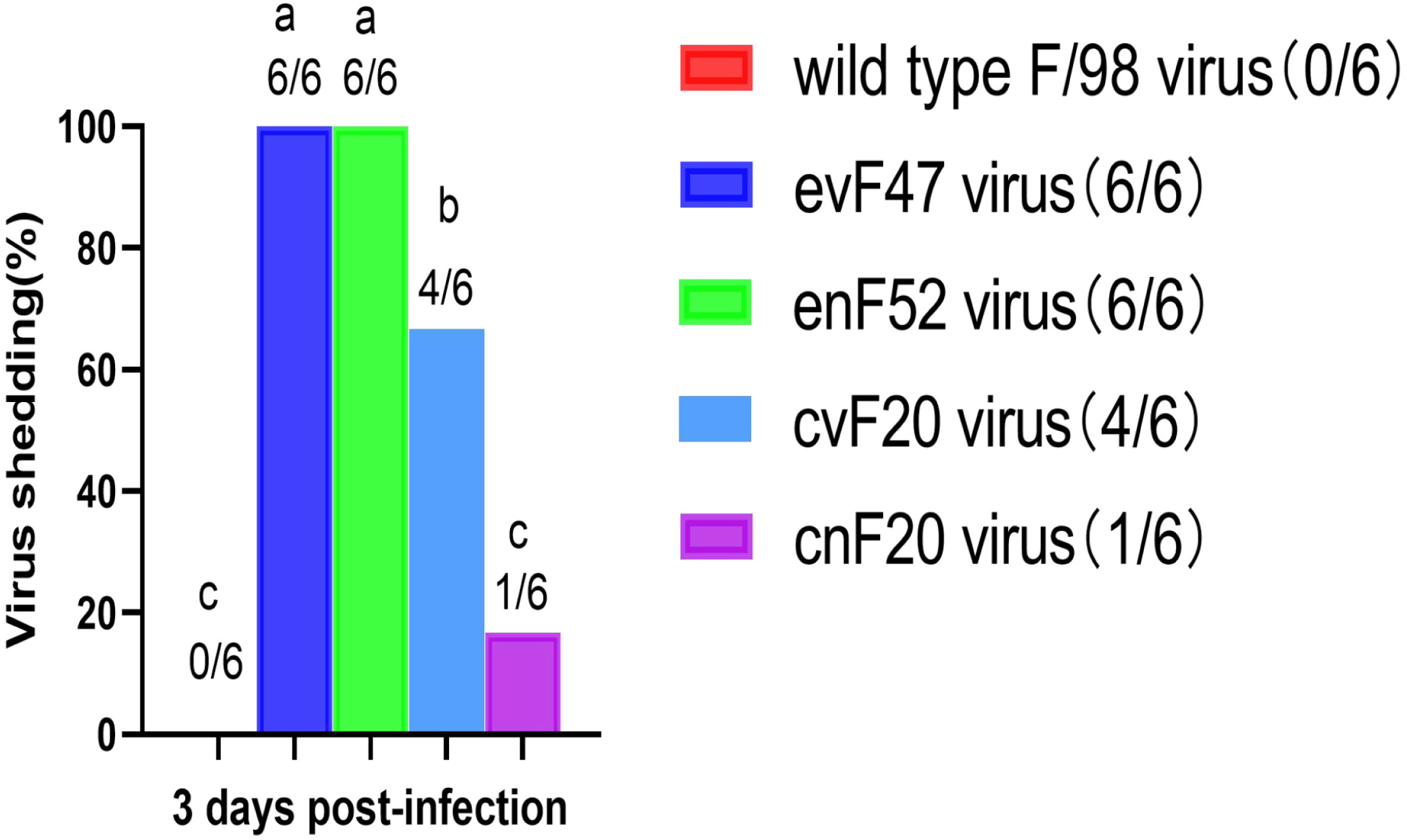
Protection efficiency of the whole inactivated F/98 vaccine against the antigen variant viruses passaged in SPF embryonated chicken eggs or in the chickens with or without homologous vaccine antibodies. The negative sample of chickens shedding virus from the swabs on 3 days post challenged with a specific virus indicated that the HA titer lower than the detection limit of 2^2^. Each group had six chickens.The appearance of the same letter means that there is no marked difference among the groups under the condition of *P* >0.05.

### HA mutations of F/98 strain under selection pressure played different roles in the antigenic variation

The passaged viruses evF47 and enF52 possessed the same mutations in HA. The S145N mutation located adjacent to the receptor-binding sites, the A198V mutation located at the receptor-binding sites, and the mutations K131R and G181E located in the HA globular domain (Figure 2A). In order to evaluate the role of these mutations, 9 recombinant viruses containing single or multiple HA mutations from the viruses evF47 and enF52 in F/98 backbone were generated, respectively, including rF/HA_K131R_, rF/HA_S145N_, rF/HA_G181E_, rF/HA_A198V_, rF/HA_348_ (K131R+S145N+G181E), rF/HA_349_ (K131R+S145N+A198V), rF/HA_389_ (K131R+ G181E +A198V), rF/HA_489_ (S145N+G181E+A198V), and rF/HA47 (K131R+S145N+G181E+A198V) (Table 1). The serum against the paternal virus F/98 in chickens was used as the reference serum to analyze the antigenicity of the recombinant viruses by HI assay. Compared with the paternal virus F/98, the mutations K131R and G181E did not affect the readouts of HI titer, the viruses rF/HA_S145N_ and rF/HA_348_ exhibited 1.67-fold lower HI titers, and the viruses rF/HA_A198V_, rF/HA_349_, rF/HA_389_, rF/HA_489_ and rF/HA47 displayed 6.67-fold lower HI titers, which were antigenically distinct from F/98 (Figure 2B). These results suggested that the mutations S145N and A198V were related to the change of the antigenicity of H9N2 virus, and the contribution of mutation A198V to the antigenic drift of F/98 strain was significantly more than that of the mutation S145N.

**Table 1.**
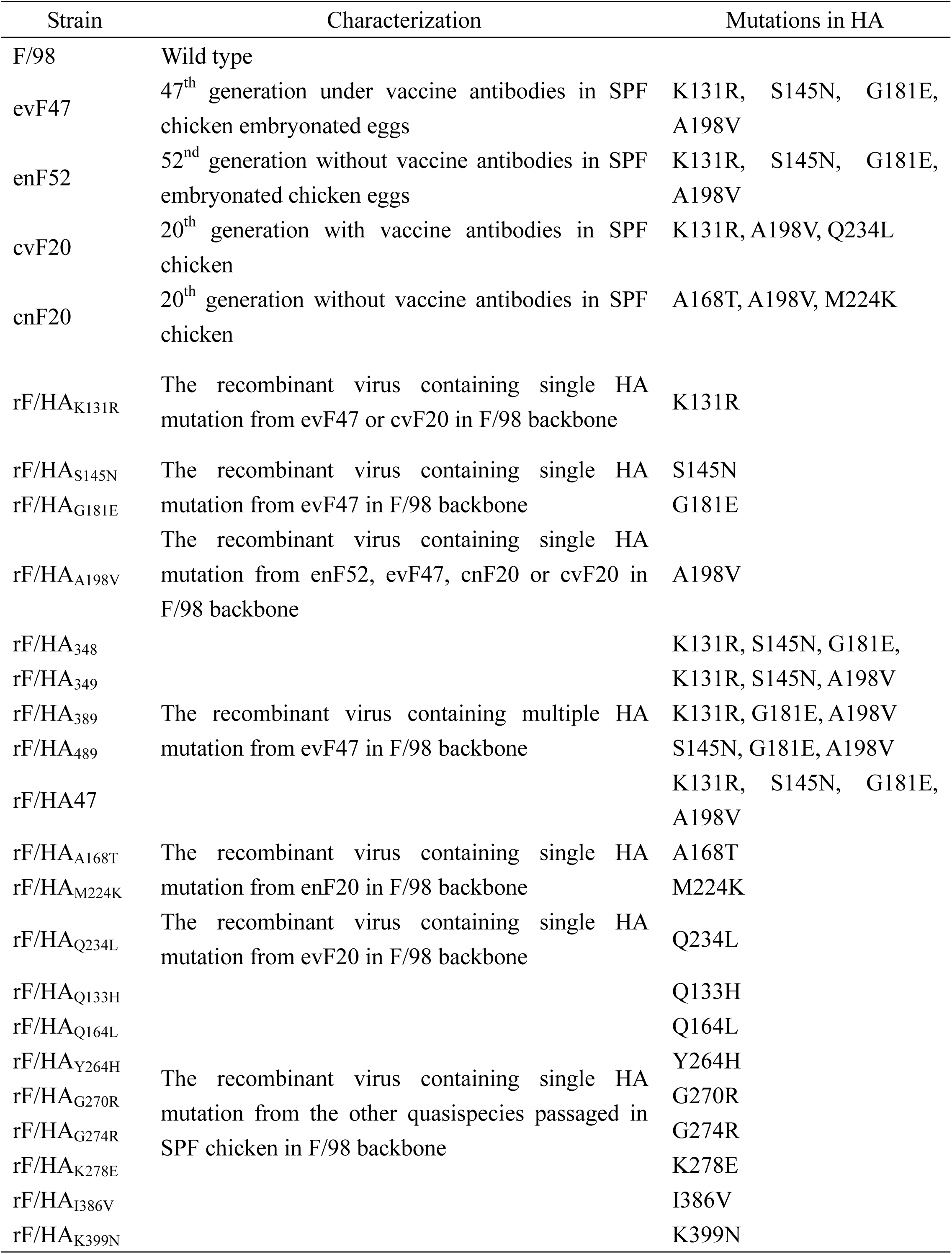
The characterization and HA mutations of the H9N2 viruses used in this study

**Figure 2.**
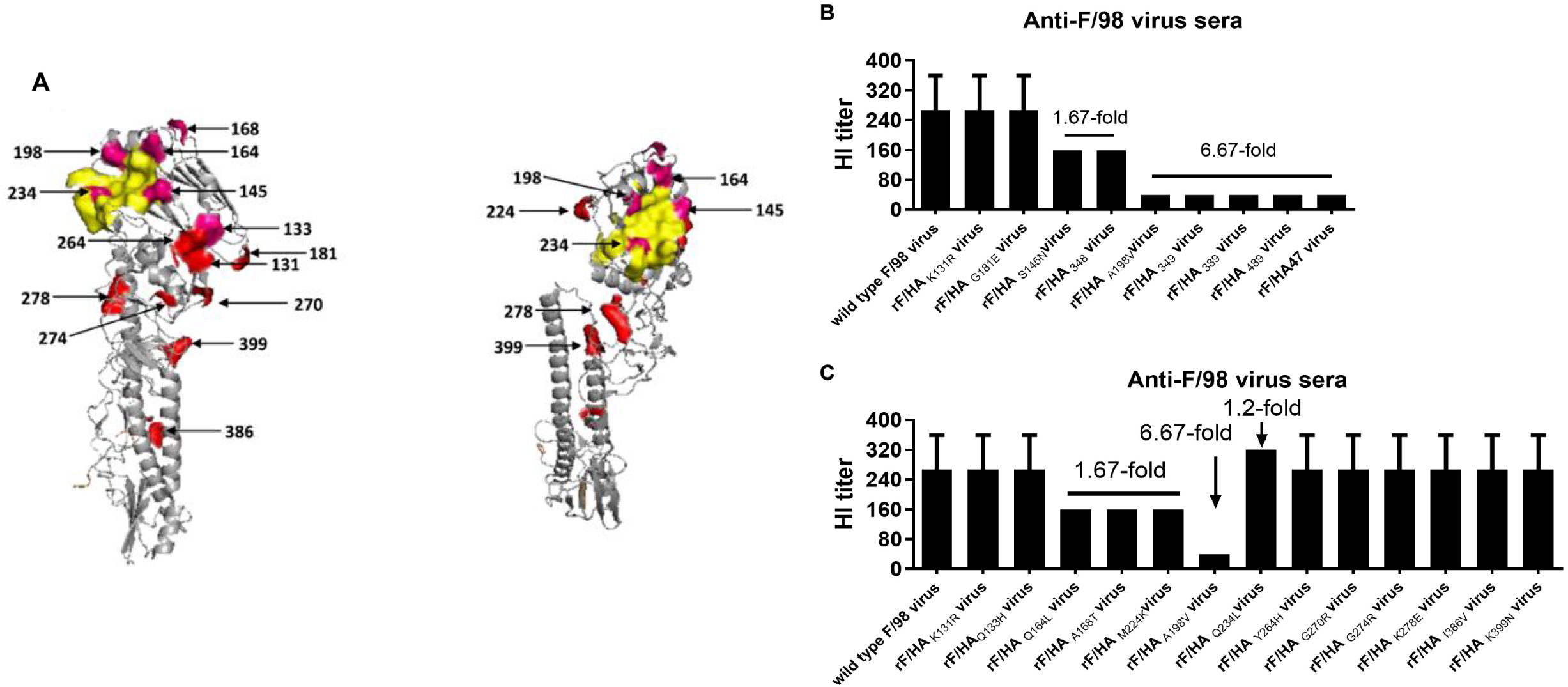
(A) The location of amino acid mutations in the three-dimensional structure of HA protein of H9N2 subtype avian influenza virus. A. Yellow color indicates the locations of the HA receptor binding sites including the positions 146-150, 109, 161, 163, 191, 198, 202, 203, and 232-237. Pink color indicates mutations on antigenic sites that have been reported. Red color indicates mutations that have not been previously reported as antigenic sites. (B) HI titers of F/98 immune sera from chickens (n=8) to each recombinant virus with single or multiple mutants in HA from the passaged viruses occurred in the 47th generation in embryonated chicken eggs under selective pressure on antibodies. (C) HI titers of F/98 immune sera from chickens (n=8) to the recombinant virus with each single mutant in HA from the quasispecies occurred in the chickens under the selection pressure of vaccine antibodies. A ≥4-fold change in HI titers of standard antiserum was considered as significant antigenic change.

In the process of F/98 strain continuously passaged in chickens, 390 quasispecies were isolated when F/98 strain was passaged for 20 generations in chickens, and 13 HA mutations were identified, including K131R, Q133H, Q164L, A168T, A198V, M224K, Q234L, Y264H, G270R, G274R, K278E, I386V and K399N (22). The mutation A198V was located at receptor-binding sites, Q164L was next to receptor-binding sites, Q234L was located in the right edge of the receptor-binding pocket, the mutations A168T and M224K were around the receptor-binding sites, and the other HA mutations except for K278E, I386V and K399N were all located in the HA globular domain (Figure. 2A). Then, 13 recombinant viruses containing single HA mutation in F/98 backbone were generated (Table 1) and were tested for their antigenicity against anti-F/98 serum using HI assay. Eight recombinant viruses, rF/HA_K131R_, rF/HA_Q133H_, rF/HA_Y264H_, rF/HA_G270R_, rF/HA_G274R_, rF/HA_K278E_, rF/HA_I386V_ and rF/HA_K399N_, had similar antigenicity to the paternal virus F/98. Three recombinant viruses, rF/HA_Q164L_, rF/HA_A168T_ and rF/HA_M224K_, exhibited 1.67-fold lower HI titers to the anti-F/98 serum. The virus rF/HA_A198V_ exhibited 6.67-fold lower HI titers (a change of more than 4-fold) to the anti-F/98 serum, which is an indication of antigenically distinct from F/98. The virus rF/HA_Q234L_ displayed 1.2-fold higher HI titer to the anti-F/98 serum (Figure. 2C). These results suggested that the mutations Q164L, A168T, A198V, M224K and Q234L were related to antigenic variation.

In conclusion, the mutations S145N, Q164L, A168T, A198V, M224K and Q234L in the HA occurred in embryonated chicken eggs or chickens with or without the selection pressure resulted in antigenic drift of F/98 strain. Among these mutations, the mutation A198V at the receptor binding site of HA significantly promoted the antigenic drift.

### HA mutations have unequal effects on the receptor binding avidity

Because the mutations S145N, Q164L, A168T, A198V, M224K and Q234L affecting the antigenicity of F/98 strain were all around the receptor binding sites of HA, we hypothesized that these mutations might alter the receptor binding avidity or the interaction between the virus and the receptor on the surface of the chicken red blood cells. The results showed that the virus cvF20, rF/HA_A198V_ and rF/HA_Q234L_ bound to chicken erythrocytes treated with 32-fold higher α2-3,6,8 neuraminidase concentrations than the F/98 strain. The virus evF47 and rF/HA47 bound to chicken red blood cells treated with 4-fold higher α2-3,6,8 neuraminidase concentrations than F/98 strain. Compared to F/98 strain, the virus rF/HA_M224K_ bound to red blood cells treated with 2-fold higher α2-3,6,8 neuraminidase concentrations, while the virus rF/HA_S145N_ bound to chicken erythrocytes less avidly (Figure 3). In order to further confirm that S145N mutation decreased receptor binding avidity in these HA mutations, a recombinant “7+1” influenza virus rF/HA47 containing HA from the virus evF47 in F/98 backbone was generated. rF/HA47 possesses K131R, S145N, G181E, and A198V mutations in HA. Four recombinant viruses in rF/HA47 backbone were generated by introducing single HA mutation R131K, N145S, E181G and V198A, respectively, namely rF/HA_489_ (S145N+G181E+A198V) (R131K in HA), rF/HA_389_ (K131R+G181E+A198V) (N145S in HA), rF/HA_349_ (K131R+S145N+A198V) (E181G in HA), and rF/HA_348_ (K131R+S145N+G181E) (V198A in HA) (Table 1). As shown in Figure 4, the receptor binding avidity of the virus rF/HA_389,_ rF/HA47 introducing the HA mutation N145S, was increased by 8-fold compared with that of the virus rF/HA47, which was consistent with the result that S145N HA mutation caused the decrease of the receptor binding avidity, and the receptor binding avidity of the virus rF/HA_348_, V198A HA mutation introduced in rF/HA47, was decreased by 4-fold compared with that of the virus rF/HA47 (Figure 3). These results indicated that the antigenic variants passaged in embryonated chicken eggs or chickens affected receptor binding avidity. The mutations A198V and Q234L significantly improved the receptor binding avidity, while the S145N mutation decreased the receptor binding avidity.

**Figure 3.**
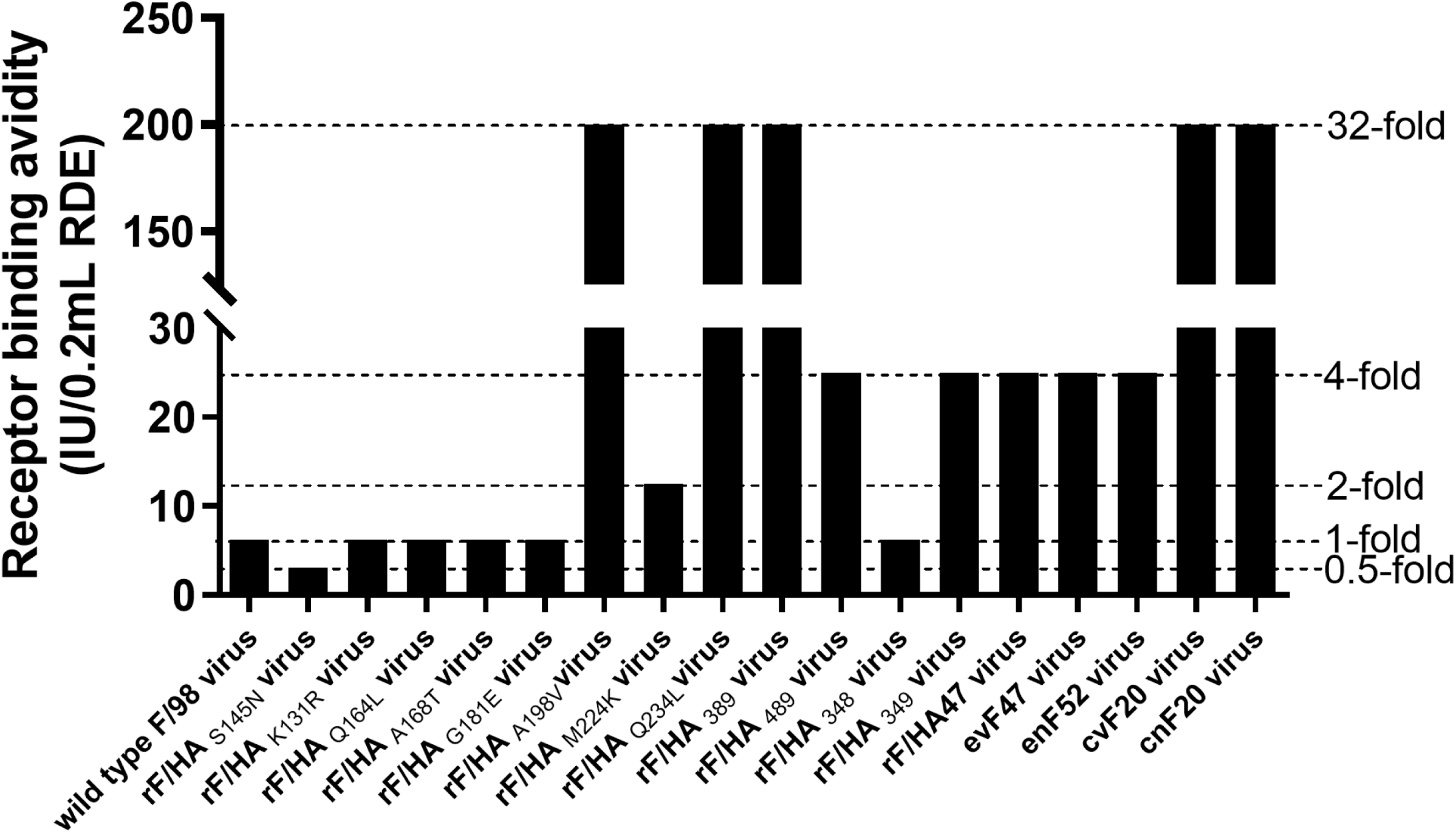
HA mutations A198V, M224K, Q234L increase receptor binding avidity of H9N2 virus F/98, whereas S145N decreases receptor binding avidity. And the evolution of F/98 strain in chicken embryos or chickens with homologous vaccine antibodies resulted in the increase of receptor binding avidity. Relative viral receptor binding avidities were determined by hemagglutination of red blood cells pretreated with increasing amounts of α2-3,6,8 neuraminidase. Data are expressed as the maximal amount of neuraminidase that allowed full agglutination. The data are representative of three independent experiments.

**Figure 4.**
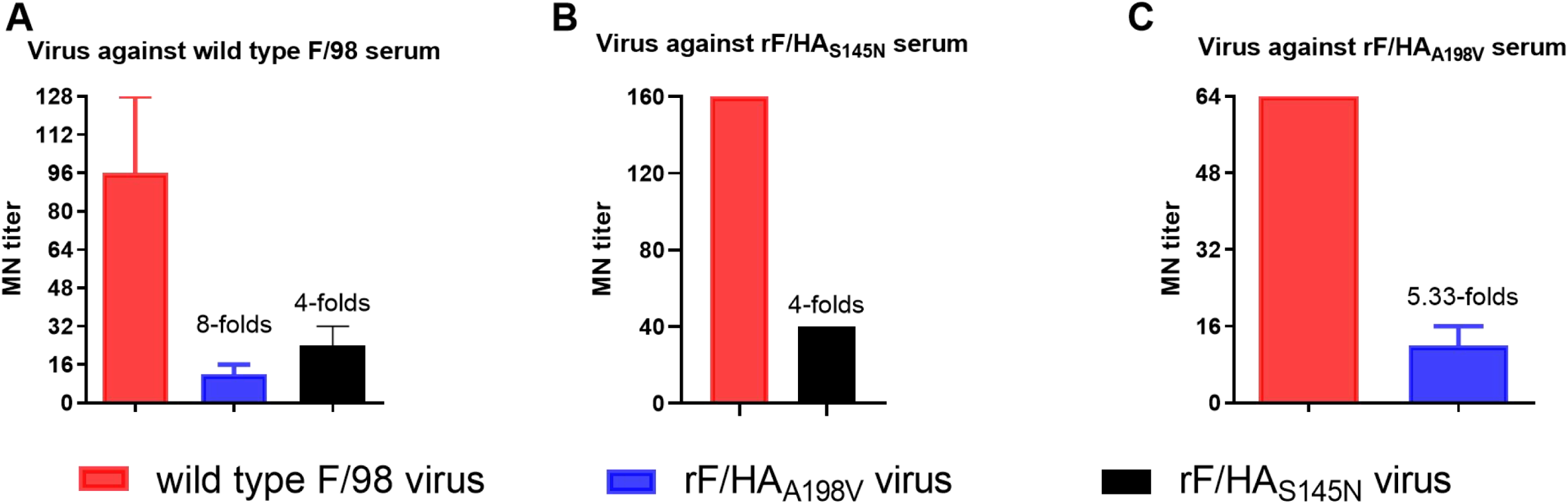
HA S145N or A198V mutation results in a decrease in MN titer. (A) MN titers of F/98 immune sera from chickens (n=8) to the viruses F/98, rF/HA_S145N_, and rF/HA_A198V_. (B) MN titers of rF/HA_S145N_ immune sera from chickens (n=8) to the viruses F/98 and rF/HA_S145N_. (C) MN titers of rF/HA_A198V_ immune sera from chickens (n=8) to the viruses F/98 and rF/HA_A198V_. A ≥4-fold change in MN titers of standard antiserum was considered as significant antigenic change.

Although the mutation S145N or A198V affected the antigenicity of H9N2 virus, the A198V mutation increased the receptor binding avidity while the S145N mutation was the opposite to the A198V mutation for receptor binding avidity, which suggested the mechanism of S145N mutation for antigenic drift might be different from that of A198V mutation. Therefore, we selected the mutations S145N and A198V to further study on the escape mechanism from selective pressure exerted by inactivated vaccine induced antibodies.

### The S145N and A196V mutations used distinct mechanisms to escape from neutralizing-antibodies

In order to study the roles of the mutations S145N and A198V in escape from neutralizing antibodies, microneutralization (MN) assay was performed, which was more sensitive than HI assay (23). In the cross-MN assay between the F/98 strain and the virus rF/HA_S145N_, the virus rF/HA_S145N_ had 4-fold lower MN titer to the anti-F/98 serum (Figure 4A), or even to the anti-rF/HA_S145N_ serum (Figure 4B). The result suggested that the S145N mutation not only caused the virus rF/HA_S145N_ escape from anti-F/98 serum, but also escape from anti-rF/HA_S145N_ serum against itself. Antibody binding ELISA confirmed that the area under the curve (AUC) of anti-F/98 serum binding to the F/98 strain was 3.2-fold higher than that of anti-F/98 serum binding to the rF/HA_S145N_ virus (*P*<0.01) (Figure 5A). The AUC of anti-rF/HA_S145N_ serum binding to the rF/HA_S145N_ virus was 2.3-fold higher than that of anti-rF/HA_S145N_ serum binding to the F/98 virus (*P*<0.001) (Figure 5B). These results revealed that the anti-F/98 serum or anti-rF/HA_S145N_ serum bound less efficiently to the rF/HA_S145N_ virus than those to the F/98 virus. These data indicated that the S145N mutation promoted virus escape from pAbs by physically preventing virus-Ab binding.

**Figure 5.**
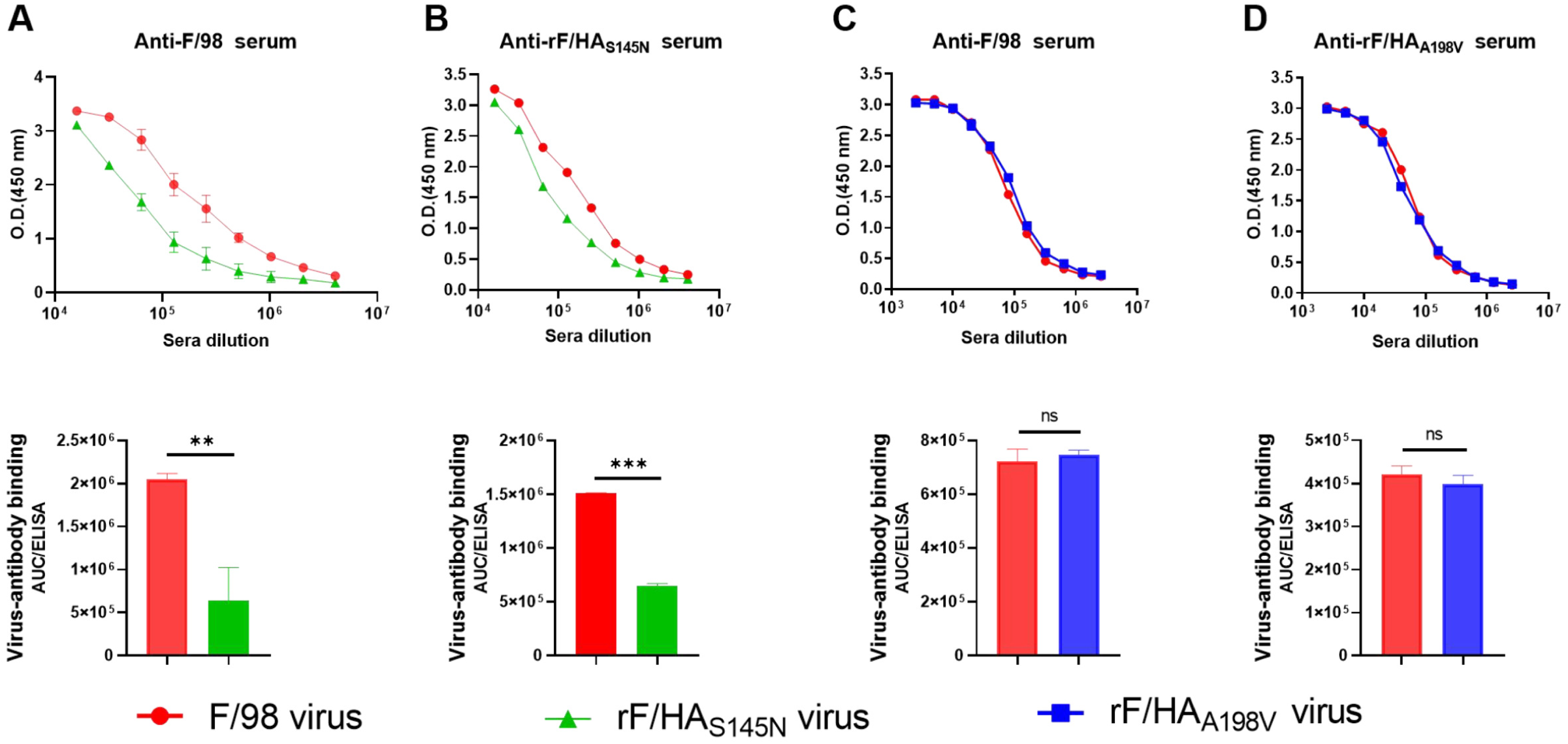
Single S145N mutation physically prevents Ab binding, whereas single A198V mutation does not affect Ab binding. Direct antibody binding to F/98 or rF/HA_A198V_ viruses were determined by ELISA using sera collected from chickens vaccinated with inactivated F/98 (A) or rF/HA_A198V_ (B). Direct antibody binding to F/98 or rF/HA_S145N_ viruses were determined by ELISA using sera collected from chickens vaccinated with inactivated F/98 (C) or rF/HA_S145N_ (D). The AUC of ELISA was calculated for virus-Abs binding by GraphPad Prism 8 software above the value of the corresponding negative control, which was performed under the same conditions. Means and SD from three independent experiments. Statistical significance was based on student’s t test (\*\**P* < 0.01; *** *P* < 0.001). O.D., optical density.

In comparation to the F/98 strain, the rF/HA_A198V_ virus exhibited 8-fold reduction of MN titers to anti-F/98 serum (Figure 4A), and 5.33-fold reduction of MN titers to homologous anti-rF/HA_A198V_ serum (Figure 4C), which suggested that the A198V mutation promoted the rF/HA_A198V_ virus escape from anti-F/98 and anti-rF/HA_A198V_ serum. Antibody binding ELISA confirmed that the anti-F/98 serum bound similarly to either the F/98 virus or the rF/HA_A198V_ virus (Figure. 5C), and anti-rF/HA_A198V_ serum also bound similarly to either the F/98 virus or the rF/HA_A198V_ virus (Figure 5D). Taken together, these data indicated that the A198V mutation promoted escape from pAb pressure by increasing viral receptor binding avidity, but not by preventing antibody binding physically.

### The mutations S145N and A198V reduced the protection efficiency of the corresponding inactivated vaccine

The anti-sera against the whole inactivated vaccine of the virus F/98 or rF/HA_S145N_ were generated in SPF chickens, and the HI assay was performed. The results showed that the average HI titer of the anti-F/98 serum against the F/98 strain was 1.83-fold higher than that against the rF/HA_S145N_ virus (*P*<0.05). Average HI titer of anti-rF/HA_S145N_ serum against the F/98 strain was 1.46-fold higher than that against the rF/HA_S145N_ virus. The HI titers of serum from the chickens vaccinated with the rF/HA_S145N_ virus were slightly lower than that from the chickens vaccinated with the F/98 virus (Figure 6A). These data suggested that the neutralizing antibody in serum induced by the inactivated vaccine of the rF/HA_S145N_ virus was slightly lower than that induced by the inactivated vaccine of the paternal virus F/98. The protection test showed that the antibody induced by the F/98 vaccine could provide 100% protection against the challenge by either the F/98 virus or the rF/HA_S145N_ virus. The antibody induced by the rF/HA_S145N_ vaccine could provide 100% protection against the challenge by the F/98 virus, while 83.3% protection against the challenge by the rF/HA_S145N_ virus (Table 2).

**Table 2.** Virus shedding from the swabs on days 3 and 5 post challenged with F/98, rF/HA_S145N_ or rF/HA_A198V_ viruses

| Group | Virus shedding# |  | Protection (%) |
| --- | --- | --- | --- |
|  | D3 | D5 |  |
| Chickens vaccinated the F/98 inactive vaccine |  |  |  |
| Challenged with F/98 virus | 0/6<sup>a</sup> | 0/6<sup>a</sup> | 100 |
| Challenged with rF/HA<sub>A198V</sub> virus | 1/6<sup>a</sup> | 0/6<sup>a</sup> | 83.3 |
| Challenged with rF/HA<sub>S145N</sub> virus | 0/6<sup>a</sup> | 0/6<sup>a</sup> | 100 |
| Chickens vaccinated the rF/HA<sub>A198V</sub> inactive vaccine |  |  |  |
| Challenged with F/98 virus | 1/6<sup>a</sup> | 0/6<sup>a</sup> | 83.3 |
| Challenged with rF/HA<sub>A198V</sub> virus | 1/6<sup>a</sup> | 0/6<sup>a</sup> | 83.3 |
| Chickens vaccinated the rF/HA<sub>S145N</sub> inactive vaccine |  |  |  |
| Challenged with F/98 virus | 0/6<sup>a</sup> | 0/6<sup>a</sup> | 100 |
| Challenged with rF/HA<sub>S145N</sub> virus | 1/6<sup>a</sup> | 0/6<sup>a</sup> | 83.3 |

**Figure 6.**
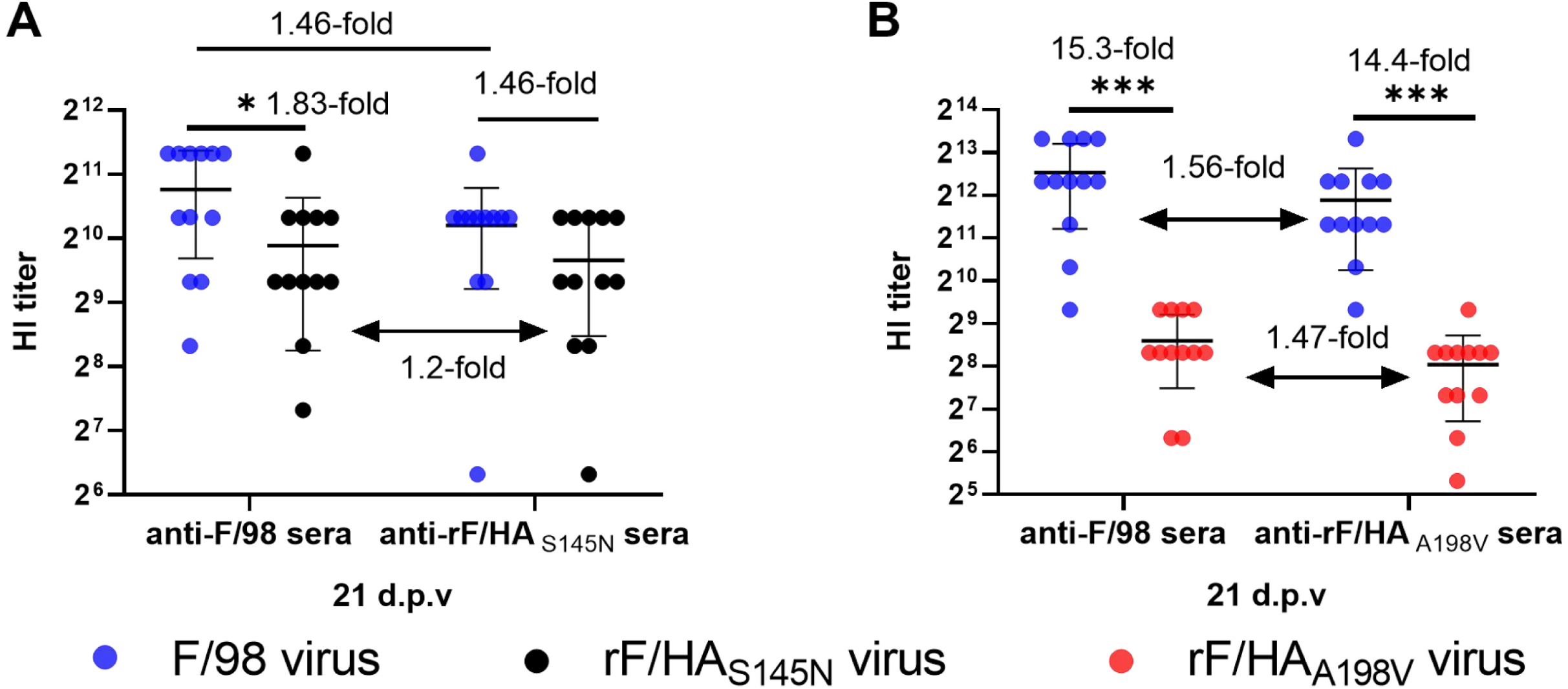
Single S145N mutation or single A198V mutation resulted in low HI reaction for chicken serum. Three-week-old SPF chickens were vaccinated once by subcutaneous injection of 0.3 mL of oil-emulsion of inactivated whole virus vaccines of the viruses F/98, rF/HA_S145N_, and rF/HA_A198V_, respectively. At 21 d.p.v., twelve chickens from each group (the F/98 vaccine group, the rF/HA_S145N_ vaccine group, and the rF/HA_A198V_ vaccine group) were bled to analyze cross-HI titers against the F/98 and rF/HA_S145N_ viruses (A), or against the F/98 and rF/HA_A198V_ viruses (B). Statistical significance was based on student’s t test (\**p* < 0.05).

Additionally, the anti-F/98 serum against the F/98 virus was 15.3-fold higher than that against the rF/HA_A198V_ virus (*P*<0.001); the anti-rF/HA_A198V_ serum against the F/98 virus was 14.4-fold higher than that against the rF/HA_A198V_ virus (*P*<0.001); the HI titer of serum from the chickens vaccinated with rF/HA_A198V_ virus were lower than that from the chickens vaccinated with F/98 virus (Figure 6B). These data demonstrated that the neutralizing-antibody in serum induced by the inactivated vaccine of the rF/HA_A198V_ virus was lower than that induced by the inactivated vaccine of the F/98 virus. The immunogenic test showed that the antibody induced by the F/98 vaccine could provide 100% protection against the challenge by the F/98 virus, and 83.3% protection against the challenge by the rF/HA_A198V_ virus; while the antibody induced by the rF/HA_A198V_ vaccine could provide 83.3% protection against the challenge by either the F/98 virus or the rF/HA_A198V_ virus (Table 2).

Taken together, these results indicated that the mutations S145N and A198V both reduced the protective efficacy of H9N2 inactivated vaccine, and promoted H9N2 virus escape from antibody-based immune response.

## Discussion

The adaptive evolution of H9N2 virus is determined by environment, genetic background and immune status, et al., resulting in introducing mutations into the viral genome of the H9N2 virus. As reported previously (21, 22), we found that the same four HA mutations occurred in the antigenic variants passaged in the 47^th^ generation under selective pressure with vaccine antibodies, or in the 52^nd^ generation under selective pressure without vaccine antibodies in SPF embryonated chicken eggs, respectively. Given that the lack of a strong immune system in embryonated chicken eggs and the presence of maternal antibodies mainly in the yolk and weakly in allantoic fluids led to the lack of sufficient immune pressure in the evolution of H9N2 virus, which ensured that the evolution of H9N2 virus in embryonated chicken eggs almost independent and free from the selection pressure, and antigenic variants occurring in embryonated chicken eggs could completely escape neutralizing antibodies induced by the paternal virus F/98 in host. Additionally, 66.67% of the antigenic variants occurring in chickens under the selection pressure of vaccine antibody could escape from antibody-based selection *in vivo*, whereas only 16.7% of the antigenic variants occurring in chickens without the selection pressure of vaccine antibody. We speculated that the adaptive evolution of H9N2 virus reflected the results of the virus-host interaction process. H9N2 virus undergoing the selection of immune pressure in chickens would develop a symbiotic relationship with an appropriate escape strategy in the process of adaptive evolution. The presence of additional selection pressure of vaccine antibody would promote the host to restrict the infection and replication of H9N2 virus, while H9N2 virus would increase the immune escape to counteract specific antibodies binding.

HA is the most important antigenic protein of H9N2 virus, which stimulates host chicken to product the HA-specific neutralizing antibodies. HA mutations in antigenic sites promoted the virus to escape from antibody-based immune responses in host. Several studies reported HA mutations that affect the antigenic variation of H9N2 virus. In these studies, HA mutations from different H9N2 antigenic variants are mainly located at or near the receptor binding sites in HA, most of which were selected with HA-specific mAbs *in vitro* (7-13). Few studies on the HA mutations selected with pAbs driving from inactivated vaccine *in vivo* and the contribution of single HA mutation in H9N2 virus to antigenic variation or immune escape were reported. Here, the contribution of 15 HA mutations, occurring *in vivo*, to antigenic variation and immune escape were studied using HA genes from antigenic variants passaged with or without selection pressure of H9N2 inactivated vaccine as our previously reported (21, 22). We found that the HA mutations S145N, Q164L, A168T, A198V, M224K, and Q234L were responsible for the antigenic variation of H9N2 virus. The A198V and Q234L mutations are at the receptor binding sites of HA. The mutations S145N, Q164L, A168T and M224K are near the receptor binding sites. The mutations S145N, Q164L and A168T have been reported when H9N2 virus were selected by either pAbs or mAbs *in vitro* (9-12). Our results also indicated that these mutations played an important role in the antigenic variation of H9N2 virus. The amino acid residues at the position 234 in HA from different H9N2 strains could be mutated under the selection pressures of either pAbs or mAbs *in vitro*, and the residue L234 was responsible to bind to human type α2,6 linked sialic acid receptors (24), suggesting the Q234L mutation occurred under selection pressure of vaccine antibodies increased the potential of F/98 strain to infect human. Although the HA mutation A198V has not been reported in antigenic mapping of HA of H9N2 virus using selection with HA-specific mAbs, about 90% of H9N2 wild viruses possess V or T at position 198 in HA (25), and the A198V mutation occurred in all of the passaged virus in this study, suggesting that the HA mutation A198V played a key role in the process of adaptive evolution of F/98 strain.

N-linked glycosylation (NLG) is a specific posttranslational modification of HA. Both NLG pattern and HA protein sequence determine the antigenic property of AIV (26). The NLG near the antigen epitope may shield the antigenic sites on the HA, causing immune escape by disturbing Abs recognition or blocking Abs binding (27, 28), and the NLG near the receptor binding sites may change its receptor-binding properties and maintain viral fitness of the receptor binding activity (29-31). For example, the mutation K144N of PR8/H1N1 virus introduced a glycosylation site in HA, and followed by the compensatory mutations D225G, N193K, or P186S that increased the receptor binding avidity (32). Naturally, the NLG at the position 145 in HA, which is present in about 10% of H9N2 isolates, is an important glycosylation site for H9N2 virus (33). In this study, the mutation S145N of the F/98 strain near the receptor binding sites of HA resulted in the addition of a glycosylation site, which shielded or interfered with the receptor sites and blocked Abs binding (9). The single S145N mutation decreased receptor binding activity, and promoted viral escape in MN or HI assays by preventing Ab binding. This finding revealed the molecular mechanism of the S145N mutation escaping from pAbs deriving from inactivated vaccine and its role in viral antigenic variation, which was not reported in the previous studies on the antigenic mutation in HA. In addition, the inactivated vaccine of the F/98 virus introduced single S145N mutation induced lower antibody level in serum, and could not provide 100% protection efficiency for its own virus rF/HA_S145N_. This is a new discovery of the immune escape strategy of the S145N mutation.

The receptor binding sites in HA of H9N2 virus include the residues at the position 109, 161, 163, 191, 198, 202 and 203, of which all are conservative except the residues at the position 198 (33). Sealy et al. investigated 55 H9N2 wild strains in Pakistan from 2014 to 2016, and found that the mutation A198V/T enhanced the receptor binding avidity of H9N2 virus (34). Herein, we demonstrated that the A198V mutation had the greatest effect on the antigenic variation of the F/98 virus among 15 HA mutations from the passaged viruses. It increased the receptor binding avidity significantly and facilitated the virus rF/HA_A198V_ to escape from the antibodies induced by the vaccine of the paternal virus F/98 or its own. Additionally, the inactivated vaccine of the rF/HA_A198V_, only introduced the A198V mutation in F/98 virus, induced lower antibody levels in serum, and could not provide 100% protection efficiency against rF/HA_A198V_. These results might explain the phenomenon that homologous vaccine antibodies could not provide acceptable protection for vaccinated chicken flocks in recent years (6).

In summary, our finding revealed that the HA mutations occurring in the selection pressure with or without vaccine antibody affected antigenic variation of H9N2 virus, including S145N, Q164L, A168T, A198V, M224K and Q234L, of which the mutation S145N or A198V significantly affected antigenic variation through different mechanisms. Further data showed that S145N substitution promoted viral escape from pAbs driving from vaccine by preventing Abs binding physically, while A198V substitution did promote H9N2 virus escape from pAbs-neutralizing reaction by enhancing the receptor binding activity. Additionally, both S145N and A198V mutations interfered with the immunogenicity of the inactivated vaccine, resulting in reduction of the protective efficiency of H9N2 inactivated vaccine, which contributed escape from the antibody-based immunity.

## Materials and methods

### Ethical compliance

The SPF chickens and chicken embryos used in this study were purchased from Nanjing Biology Medical Factory, Qian Yuan-hao Biological Co, Ltd.. Procedures involving the care and use of animals were approved by the Jiangsu Administrative Committee for Laboratory Animals (permission number SYXK 2016-0020) and performed in accordance with the Jiangsu Laboratory Animal Welfare and Ethics guidelines of the Jiangsu Administrative Committee of Laboratory Animals.

### Viruses and cells

The H9N2 virus F/98 was isolated in Shanghai in 1998, stored at - 70 °C at the Animal Infectious Disease Laboratory, School of Veterinary Medicine, Yangzhou University. The GenBank accession numbers of the sequence of the F/98 strain are AY253750-AY253756 and AF461532 (35). Human embryonic kidney cells (293T) and Madin-Darby canine kidney (MDCK) cells, purchased from ATCC (Manassas, VA, USA), were maintained in Dulbecco’s modified Eagle’s medium (DMEM) (Sigma, St. Louis, MO, USA) supplemented with 10% fetal calf serum (Hyclone, South Logan, UT, USA) and were incubated at 37 °C with 5% CO_2_.

### Generation of H9N2 AIVs by reverse genetics

The primers, synthesized by Tsingke Biological Technology (Nanjing, China), used to amplify the DNA sequence to add the single mutation in HA protein of F/98 virus were designed using Primer 5.0 software (Primer-E Ltd., Plymouth, UK) based on the HA gene sequence of the F/98 H9N2 avian influenza virus. The full-length HA genes containing the single mutation was amplified by PCR, and inserted into a transcriptional/expression vector pHW2000 by using ClonExpress II One Step Cloning Kit (Vazyme Biotech Co., Ltd., Nanjing, China), resulting in the plasmids pHW204-HA145N. Seven dual-promoter plasmids, including pHW201-PB2, pHW202-PB1, pHW203-PA, pHW205-NP, pHW206-NA, pHW207-M, and pHW208-NS, from the F/98 virus strain possessing S145 HA, were stocked in our lab at -70 °C (36). The recombinant viruses were rescued by transfection in the 293T cell as previously described (37). Briefly, a total weight of 2.4 ng of the eight plasmids mixture with a rate of 1:1 was mixed with 100 µL Opti-MEM medium (GIBCO, BRL, Grand Island, USA). Next, 7 µL of PolyFect transfection reagent (QIAGEN, Duesseldorf, Germany) was added. The samples were incubated at room temperature for 10 min and then added to the 70-80% confluent monolayers of 293T cell in 24-well plates. After incubation at 37°C with 5% CO_2_ for 6 h, 2 µg/mL of TPCK-trypsin (Sigma, St. Louis, MO, USA) was added to the wells. Thirty hours after transfection, the supernatants were harvested and inoculated into 10-day-old SPF embryonated chicken eggs for virus propagation. The rescued virus was analyzed with a hemagglutinin assay, and the HA genes from the rescued virus was sequenced by Tsingke Biological Technology (Nanjing, China) to confirm the accuracy of the designed mutation.

### Determination of the 50% tissue cell infectious dose (TCID_50_)

The TCID_50_ assay was performed as we described previously (37). Briefly, the viruses were diluted in DMEM without serum to a concentration of 10^−1^ to 10^−11^ and then added to MDCK cells in 96-well plates, respectively. After incubation at 37 °C with 5% CO_2_ for 1 h, the supernatants were removed. The plates were washed twice with PBS, and then 100 µL of DMEM was added to each well. After incubation at 37°C with 5% CO_2_ for 72 h, the HA titers of the cell supernatants were analyzed. The virus titers were calculated according to the Reed-Muench formula.

### Anti-sera

As described previously (25), six three-week-old SPF chickens were immunized twice by subcutaneous injection of 0.3 mL of oil-emulsion of inactivated whole virus vaccines of the viruses F/98, rF/HA_S145N_ and rF/HA_A198V_, which were inactivated by adding 0.2% formalin (*v*/v) for 24 h at 37 °C, respectively. The antisera were collected and pooled from the vaccinated SPF chickens at three weeks after the vaccination.

### Hemagglutinin-inhibition (HI) assay and Microneutralization (MN) assay

Antisera were treated with cholera filtrate (Sigma-Aldrich, St. Louis, MO, USA) to remove nonspecific hemagglutination inhibitors before HI assay. HI assay was performed using 4 hemagglutination units (HAU) of H9N2 and 1% (*v*/v) chicken erythrocytes as we described previously (37).

MN assay was performed as previously described (38). Briefly, the sera were serially diluted with 100 TCID_50_ virus and incubated at 37 °C with 5% CO_2_ for 1 h. The serum-virus mixtures were added to MDCK cells and incubated for 1 h. After incubation, the serum-virus mixtures were removed. Serum-free DMEM containing 2 µg/mL TPCK-trypsin was added to each cell and incubated at 37°C and 5% CO_2_. After 72 h of incubation, culture supernatant was mixed with equal volume of 1% (*v*/v) chicken erythrocytes to confirm the existence of hemagglutination by virus. The MN titer was defined as the highest dilution of serum with absence of hemagglutination.

### Enzyme-linked immunosorbent assay (ELISA)

ELISA assay was performed as we described previously (37). Sucrose gradient-purified viruses were diluted in PBS and added to Nunc-Immuno MaxiSop 96-well plates (Corning, NY, USA) at 16 HAU per well. After incubation overnight at 4°C, samples in wells were blocked with PBS-nonfat dry milk. Antisera against the F/98 virus, the rF/HA_S145N_ virus or the rF/HA_A198V_ virus in chickens were then added in serial twofold dilutions with PBS containing 0.05% Tween 20, respectively, and incubated for 3 h at 37 °C. After washing, goat anti chicken horseradish peroxidase antibody (Abcam, Cambridge, MA) was added and allowed to incubate for 1.5 h at 37 °C. After washing, TMB (3,3′,5,5′ Tetramethylbenzidine) (Sigma, St. Louis, MO, USA) substrate was added, and the reaction was stopped by adding H_2_SO_4_. Absorbance was recorded at 450 nm using an automated ELISA plate reader (model EL311SX; Biotek, Winooski, VT). The area under curve (AUC) of either virus was assessed for virus-Abs binding with GraphPad Prism 8 software (San Diego, CA) above that of the corresponding negative control.

### Receptor binding assay

Receptor binding assay was performed as previously described (39). Briefly, the chicken erythrocytes were pretreated with different amounts of α2-3,6,8 neuraminidase (New England Biolabs, Beverly, MA, USA) for 1 h at 37 °C. The chicken erythrocytes were washed with PBS and added (as 1% (*v*/v) solutions) to 4 HAU of each virus (as determined using nontreated chicken erythrocytes). Agglutination was measured after incubation for 1 hour. Virus with higher receptor binding avidity is able to bind to chicken erythrocytes that are treated with high amounts of α2-3,6,8 neuraminidase.

### Chicken experiments

(i) A total of 30 three-week-old SPF chickens vaccinated with the whole inactivated F/98 virus were divided into five groups: the F/98 challenge group, the evF47 challenge group, the enF52 challenge group, the cvF20 challenge group, and the cnF20 challenge group. Each group has 6 chickens. At day 21 post-vaccination, each group was challenged intranasally and intratracheally with 10^6^ EID_50_ of the corresponding virus. ii) A total of 42 three-week-old SPF chickens were divided into three groups: the F/98 vaccine group including 18 chickens, the rF/HA_S145N_ vaccine group including 12 chickens, and the rF/HA_A198V_ vaccine group including 12 chickens, which were immunized with the emulsion vaccine of the F/98 virus, the rF/HA_S145N_ virus, and the rF/HA_A198V_ virus, respectively. At day 21 post-vaccination, chickens were bled from the wing vein for sera, and HI reactions against the F/98 virus, the rF/HA_S145N_ virus or the rF/HA_A198V_ virus was performed, respectively. Then, six chickens from each group were challenged intranasally and intratracheally with 10^6^ EID_50_ of the F/98 virus, the rF/HA_S145N_ virus or the rF/HA_A198V_ virus. Chickens were monitored daily for morbidity and mortality after challenge. At days 3 and 5 post-challenge, tracheal and cloacal swabs from challenged chickens were collected in 1 mL of PBS containing antibiotics. After one freeze-thaw cycle, the swabs were centrifuged at 3000 rpm for 10 min. A 0.2 mL supernatant was taken to inoculate 10-day-old SPF chicken eggs. Viral shedding in the trachea and cloacal was evaluated via HA titers of the allantoic cavity of SPF chicken eggs at day 5 post-inoculated according to the standard of HA≥2^3^.

### Statistics analysis

Data were shown as the mean ± *SD* for all assays. The Student’s t test analysis was used to compare between different groups and analyzed with GraphPad Prism 8 software. Differences were considered statistically significant when a *P* value was <0.05.

